# PV+ optogenetic stimulations at specific frequencies in specific brain regions can restore navigational flexibility in an acute MK801 mouse model of schizophrenia

**DOI:** 10.1101/2024.06.28.601158

**Authors:** Enrico Patrono, Daniela Černotová, Jan Svoboda, Aleš Stuchlík

**Affiliations:** Institute of Physiology of the Czech Academy of Sciences, Videnska, 1830, 14200 Prague 4, Czech Republic; Center for Advanced Behavioral Research (CABR), School of Psychology, University of New York in Prague (UNYP), Londynska 41, 12000, Prague 2, Czech Republic; Third Faculty of Medicine, Charles University, Ruska 87, 100 00 Prague 10, Czech Republic

**Keywords:** schizophrenia, navigational flexibility, NMDARs/GABA ratio, medial prefrontal cortex/ventral hippocampus, gamma/theta waves, in vivo optogenetics

## Abstract

Impairments of decision-making and behavioral flexibility in schizophrenia (SCZ) are currently the most investigated features. One convincing hypothesis explaining this cognitive impairment is the excitatory/inhibitory (E/I) ratio imbalance in brain regions such as the medial prefrontal cortex (mPFC) and the ventral hippocampus (vHPC). An increased GLUergic excitatory activity and a decreased GABAergic inhibitory activity induces an mPFC-vHPC γ/θ band desynchronization in many tasks testing behavioral flexibility. However, these tasks were carried out using “perceptual” decision-making/flexibility but not navigational decision-making/flexibility. Our study addressed the role of frequency-specific optogenetic stimulations of GABAergic parvalbumin-positive (PV+) interneurons in mPFC (50Hz, γ-like) and vHPC (10Hz, θ-like) in an acute-MK801 mouse model of navigational inflexibility. We used the active place avoidance task on a rotating arena. Results showed that frequency-specific optogenetic stimulations of mPFC or vHPC acted differently in restoring navigational flexibility, advancing our knowledge of the pivotal role of PV+ activity in SCZ-like navigational decision-making/flexibility.

## Introduction

In schizophrenia (SCZ), impaired executive functions are crucial symptoms and essential markers of SCZ onset [1]. They can significantly impact an individual’s decision-making ability and adaptation to changing circumstances. Recent reports using SCZ rodent models showed poor *behavioral flexibility* in “perceptual” decision-making tasks [2, 3]. *Behavioral flexibility* refers to the ability to reverse or switch a behavior to solve a task [4]. And *decision-making* is a complex ability that requires the evaluation of options, the formation of a preference, the execution of an action, and the processing of the consequences [5]. Specifically, “perceptual” decision-making can be explained as a flexible choice based on task rules and the subject’s perceptions, waiting to receive sensory stimuli and making the correct choice [2, 3]. On the other hand, spatial navigation is another crucial and possibly compromised behavior in SCZ. It is the ability to navigate flexibly across detours and shortcuts, finding a new path when a familiar route is blocked [6].

Interestingly, most of the behavioral paradigms investigating flexibility in decision-making did not involve spatial navigation [7–8], and most of those investigating spatial navigation did not involve decision-making [9–10]. Although perceptual decision-making tasks may represent one way to measure behavioral flexibility and executive functions, recent human and animal studies focused on cognitive maps and somatosensory circuits [11–12]. Such studies investigated the encoding of spatial variables, such as the animal’s location, heading direction, running velocity, and spatial memory. Cortical and subcortical areas have been identified as having a causal role in flexible decisions (medial prefrontal cortex and ventral hippocampus, in particular) [8, 12–13], but it remains unclear whether these areas are involved in flexible decisions during navigation. Here, we sought to determine how specific optogenetic manipulations influence navigational flexibility in response to sensory-based risk cues, thus seeking to evaluate the role of prefrontal-hippocampal crosstalk in flexible decisions during spatial navigation.

In the last decade, research in SCZ-related physiological pathways of cognitive impairments highlighted the critical role of N-methyl-D-aspartate subtype’s glutamatergic receptors (NMDARs) hypofunction in parvalbumin-positive (PV+) GABAergic interneurons with observed γ-wave desynchronization in SCZ animal models [14–16]. A convincing hypothesis explaining the cognitive impairment was connected to excitatory/inhibitory (E/I) ratio imbalance. The E/I balance refers to the balanced state of singular entities’ overall excitatory and inhibitory levels at single-cell and global circuit levels [17]. Sohal and Rubenstein [18] have clarified the multidimensional concept of this balance, stating that excitatory and inhibitory signals originate from multiple sources and can act on different targets. This condition raised discussions on enhancing the PV+ inhibitory activity to restore the E/I balance [19], resynchronizing γ-wave oscillations [20], and potentially restoring cognitive abilities [21]. Previously, we found that MK801-induced behavioral inflexibility during a perceptual decision-making task was rescued using frequency-specific optogenetic stimulations of PV+ in the medial prefrontal cortex (mPFC, 50Hz) and the ventral hippocampus (vHPC, 10Hz) [2]. Notably, acute MK801 systemic injections induce acute NMDARs hypofunction, resulting in increased cortical activity and decreased GABAergic inhibitory activity (E/I imbalance) [22]. Consequently, the E/I imbalance reduces mPFC-γ and vHPC-θ oscillations, leading to an mPFC-γ/vHPC-θ bands desynchronization [18, 23]. Furthermore, PV+ GABAergic interneurons can regulate the firing rate, spike timing, and synchronization of excitatory cells [24–25]. Finally, the enhancement of γ-/θ-waves coupling may reflect a compensatory mechanism to level the increasing difficulty of cognitive tasks [23].

Optogenetic manipulations and virtual mazes have been used to systematically screen the contributions of a wide range of cortical areas (visual cortices, retro splenial cortices, dorsal hippocampi) [11–12], brilliantly showing a complex interaction between visual and memory brain areas. However, to our knowledge, whether an altered navigational flexibility in mice can be restored by using optogenetic stimulations of PV+ interneurons at specific frequencies in the mPFC and vHPC has not yet been examined.

Here, we performed three experiments using a modified version of the active place avoidance (APA) task on a rotating arena to study *navigational flexibility*, namely reversal spatial memory and navigation set-shifting, in an acute MK801 mouse model of SCZ. Firstly, we delivered frequency-specific optogenetic stimulations to PV+ interneurons in mPFC (50Hz, γ-like) and vHPC (10Hz, θ-like) to rescue flexible navigation during the navigational flexibility sessions. Secondly, we compared the “optogenetic rescue” with control groups receiving only MK801 or NaCl i.p. injections to evaluate the therapeutical potential of the PV+ optogenetic stimulations. Thirdly, we compared the previous PV+ optogenetic stimulations-MK801 i.p. pairings with PV+ optogenetic stimulations-NaCl i.p. pairings to rule out any unspecific MK801 i.p. effect. Further, using immunohistochemical (IHC) assays, we evaluated whether mpFC and vHPC PV+ interneurons were adequately activated by measuring the levels of colocalization of c-Fos protein on the PV+ interneurons. Detailed methods are described in supplementary materials (Materials and Methods, Suppl. Mat.).

We showed that frequency-specific optogenetic stimulations of PV+ interneurons in mPFC or vHPC differently restored spatial reversal learning and navigational set-shifting. The γ-like frequency in mpFC partly restored the spatial reversal memory, and significantly restored the navigation set-shifting. Conversely, the θ-like frequency in vHPC only significantly rescued spatial reversal learning, while navigational set-shifting was partly restored. IHC assays showed high levels of c-Fos/PV+ colocalization in the “optogenetic-sensitive” groups compared to controls, confirming that frequency-specific optogenetic stimulations activated region-specific PV+ interneurons properly, thus confirming the behavioral results. Our results suggest that despite the consensus that navigation abilities are of a hippocampal domain, behavioral flexibility during various navigational strategies is more under the control of prefrontal regions.

## Results

We evaluated the behavioral results from the APA task and the effectiveness of the local (mPFC and vHPC) optogenetic stimulations of PV+ interneurons. The results are sectioned and shown in accordance with the three experiments.

The behavioral analysis was performed considering the following parameters: total distance moved, the number of entrances to the target sector, the time to enter in the target sector, and the time spent in the target sector. These parameters were considered regarding the previously reported behavioral effects of MK801 [35–36].

In the Experiment 1, we measured the behavioral parameters in the acquisition, reversal, and set-shifting sessions. Following, we used the same behavioral measurements and compared the last day of acquisition (Acq3) with reversal days (Rev1->Rev3), and the last day of reversal (Rev3) and set-shifting days (Set1->Set3) to evaluate the ability of optogenetic stimulations to restore navigational flexibility.

In the Experiment 2, we evaluated whether the frequency-specific optogenetic stimulations in mPFC and vHPC restored aspects of navigational flexibility that could resemble those of control groups (NaCl i.p.). Therefore, we compared the previous “optogenetic-sensitive” groups (WT and PV-Cre, both mPFC and vHPC) to groups mice receiving only NaCl or MK801 i.p. injections, during the reversal and the set-shifting sessions.

In the Experiment 3, we sought to determine whether MK801 i.p. injections *per se* were affecting the behavioral outcomes; and whether optogenetic stimulations were properly effective in counteracting the MK801 administration. To this aim, we compared the initial “optogenetic-sensitive” groups paired with acute MK801 i.p. injections to other “optogenetic-sensitive” groups of mice receiving NaCl i.p. injections, during reversal and set-shifting sessions.

We divided the analysis into three main groups comparisons, according to the three experiments. Table 1 summarizes the analysis strategy (Tab 1). A detailed description of the results with all the reported values is in the Supplementary Materials (Detailed results, Suppl. Mat.).

**Tab 1.** Summary of the measured behavioral parameters. Numbers in parenthesis represent the analysis groups: 1) comparisons between mPFC and vHPC groups (WT vs. PV-Cre); 2) comparisons between the previous groups and pharmacological control groups (NaCl vs. MK801) during reversal and set-shifting sessions; and 3) comparisons between mPFC and vHPC groups (WT vs. PV-Cre, with MK801 i.p.) and their control groups (WT vs. PV-Cre, with NaCl i.p.).

| PARAMETERS | BEHAVIORAL SESSIONS |  |  |
| --- | --- | --- | --- |
|  | ACQUISITION | REVERSAL | SET-SHIFTING |
| TOTAL DISTANCE MOVED | Acq1->Acq3 (1) | Rev1->Rev3 (1, 2, 3) | Set1->Set3 (1, 2, 3) |
|  |  | Acq3-Rev1->Rev3 (1) | Rev3-Set1->Set3 (1) |
| N° OF ENTRANCES | Acq1->Acq3 (1) | Rev1->Rev3 (1, 2, 3) | Set1->Set3 (1, 2, 3) |
|  |  | Acq3-Rev1->Rev3 (1) | Rev3-Set1->Set3 (1) |
| LATENCY FIRST ENTRANCE | Acq1->Acq3 (1) | Rev1->Rev3 (1, 2, 3) | Set1->Set3 (1, 2, 3) |
|  |  | Acq3-Rev1->Rev3 (1) | Rev3-Set1->Set3 (1) |
| TIME IN THE TARGET SECTOR | Acq1->Acq3 (1) | Rev1->Rev3 (1, 2, 3) | Set1->Set3 (1, 2, 3) |
|  |  | Acq3-Rev1->Rev3 (1) | Rev3-Set1->Set3 (1) |

**Table 2:** Summary of the two-ways RM ANOVAs of acquisition, reversal and set-shifting sessions. Significant values are in bold.

| *Acquisition (Acq1->Acq3)* |  |  |  |
| --- | --- | --- | --- |
| *Behavioral Parameters* | *Groups Effect* | *Days Effect* | *Interaction Effect* |
| *Tot distance moved* | **F (3, 28) = 6.441 P=0.0019** | F (1.962, 54.93) = 1.974<br>P=0.1495 | F (6, 56) = 1.365<br>P=0.2448 |
| *N° of entrances* | F (3, 28) = 0.3008 P=0.8245 | **F (1.959, 54.84) = 29.92**<br>**P<0.0001** | F (6, 56) = 1.274<br>P=0.2841 |
| *Latency to first entrance* | $F(3, 28) = 2.059$ $P=0.1284$ | $F(1.675, 46.91) = 24.71$ $P<0.0001$ | $F(6, 56) = 2.552$ $P=0.0296$ |
| *Time spent in target sector* | $F(3, 28) = 7.214$ $P=0.0010$ | $F(1.745, 48.87) = 8.542$ $P=0.0011$ | $F(6, 56) = 0.03000$ $P=0.9999$ |
| **Reversal (Rev1->Rev3)** |  |  |  |
|  | **Groups Effect** | **Days Effect** | **Interaction Effect** |
| *Tot distance moved* | $F(3, 28) = 11.38$ $P<0.0001$ | $F(1.847, 51.71) = 3.191$ $P=0.0531$ | $F(6, 56) = 1.828$ $P=0.1101$ |
| *N° of entrances* | $F(3, 28) = 1.982$ $P=0.1396$ | $F(1.569, 43.93) = 7.097$ $P=0.0041$ | $F(6, 56) = 3.615$ $P=0.0042$ |
| *Latency to first entrance* | $F(3, 28) = 15.54$ $P<0.0001$ | $F(1.968, 55.12) = 2.788$ $P=0.0711$ | $F(6, 56) = 1.598$ $P=0.1648$ |
| *Time spent in target sector* | $F(3, 28) = 10.38$ $P<0.0001$ | $F(1.966, 55.04) = 16.16$ $P<0.0001$ | $F(6, 56) = 1.932$ $P=0.0914$ |
| **Set-Shifting (Set1->Set3)** |  |  |  |
|  | **Groups Effect** | **Days Effect** | **Interaction Effect** |
| *Tot distance moved* | $F(3, 28) = 11.56$ $P<0.0001$ | $F(1.887, 52.83) = 0.2376$ $P=0.7766$ | $F(6, 56) = 1.039$ $P=0.4102$ |
| *N° of entrances* | $F(3, 28) = 13.67$ $P<0.0001$ | $F(1.628, 45.59) = 2.429$ $P=0.1093$ | $F(6, 56) = 1.639$ $P=0.1536$ |
| *Latency to first entrance* | $F(3, 28) = 7.089$ $P=0.0011$ | $F(1.912, 53.53) = 3.846$ $P=0.0292$ | $F(6, 56) = 3.339$ $P=0.0070$ |
| *Time spent in target sector* | $F(3, 28) = 4.521$ $P=0.0105$ | $F(1.885, 52.77) = 5.515$ $P=0.0076$ | $F(6, 56) = 4.805$ $P=0.0005$ |

### Comparisons between mPFC and vHPC: the role of optogenetic stimulation

At first, we evaluated if the animal groups learned the initial place avoidance task equally. A two-way RM ANOVA revealed an effect of the days (Acq1, Acq2, Acq3) in the n° of entrances, in the latency to enter the target sector, and in the time spent in the target sector, thus suggesting that mice of all groups learned the initial place avoidance task equally.

Subsequently, we evaluated whether the mice could reverse the previous spatial acquisition based on the RF rule. A two-way RM ANOVA revealed an effect of the days (Rev1, Rev2, Rev3) in the time spent in the target sector, suggesting that optogenetic stimulations in both mPFC and vHPC at specific frequencies helped to reverse the previously acquired target sector. Furthermore, an effect of the groups was found in the total distance moved and in the latency to the target sector. Finally, multiple comparisons confirmed the restoring ability of the optogenetic stimulations in mPFC (50Hz) and in vHPC (10Hz) compared to the WT groups.

Following, we evaluated whether the mice could flexibly shift the RF-to-AF rule and navigate flexibly in the arena. A two-way RM ANOVA revealed an effect of the groups in the total distance moved, in the n° of entrances, in the latency to enter the target sector, and in the time spent in target sector, suggesting that optogenetic stimulations in the mPFC at 50Hz significantly helped in flexibly shifting the RF-to AF rule compared to the WT groups, and significantly helped the flexibility also compared to the vHPC PV-Cre group. Moreover, an effect of the days was found in the latency to enter the target sector and in the time spent in the target sector, suggesting an ability of the optogenetic stimulations to induce a new spatial acquisition pattern. Finally, multiple comparisons suggest an overall ability to switch from reversal spatial memory to navigational flexibility when mPFC and vHPC are stimulated at specific frequencies.

However, the previous analysis only established that optogenetic stimulations in both mPFC and vHPC faced the MK801 challenge in reversal spatial memory and navigational set-shifting during the days. These results did not fully clarify if PV+ optogenetic stimulations in mPFC and vHPC could rescue those executive functions. To verify this, we compared the last day of acquisition (Acq3) with the reversal (Rev1, Rev2, Rev3) and the last day of reversal (Rev3) with the set-shifting (Set1, Set2, Set3). Results are shown in Table 3 (Tab. 3). Two-way RM ANOVA comparing Acq3 with reversal revealed a significant effect of the sessions in the n° of entrances, in the latency to enter the target sector, and the time spent in the target sector. Furtherly, a significant effect of the groups was found in the total distance moved, and in the time spent in the target sector. Multiple comparisons showed that *γ*-like 50Hz optogenetic stimulations in the mPFC had an immediate and generally lasting effect in reversing the spatial memory of the to-be-avoided sector compared to the WT and the PV-Cre-vHPC groups. Of note, the vHPC-PV-Cre group showed better ability to reverse the time spent in the target sector than the mPFC-Pv-Cre group, suggesting that *θ*-like 10Hz optogenetic stimulations rescued the flexibility to reverse the previous spatial memory better than *γ*-like 50Hz in mPFC.

**Table 3.** Summary of the two-ways RM ANOVAs of comparisons between the last day of acquisition and reversal, and the last day of reversal and set-shifting sessions. Significant values are in bold.

|  |  |  |  |
| --- | --- | --- | --- |
| **Reversal (Acq3 - Rev1->Rev3)** |  |  |  |
| **Behavioral Parameters** | **Groups Effect** | **Sessions Effect** | **Interaction Effect** |
| *Tot distance moved* | **F (3, 28) = 11.77**<br>**P<0.0001** | **F (2.732, 76.49) = 3.352**<br>**P=0.0268** | **F (9, 84) = 3.738**<br>**P=0.0005** |
| *N° of entrances* | F (3, 28) = 1.598<br>P=0.2122 | **F (2.160, 60.49) = 29.98**<br>**P<0.0001** | **F (9, 84) = 3.280**<br>**P=0.0018** |
| *Latency to first entrance* | **F (3, 28) = 18.00**<br>**P<0.0001** | **F (1.431, 40.07) = 58.59**<br>**P<0.0001** | **F (9, 84) = 4.003**<br>**P=0.0003** |
| *Time spent in target sector* | **F (3, 28) = 16.49**<br>**P<0.0001** | **F (2.795, 78.27) = 41.17**<br>**P<0.0001** | **F (9, 84) = 3.073**<br>**P=0.0031** |
| ***Set-Shifting (Rev 3 - Set1->Set3)*** |  |  |  |
|  | *Groups Effect* | *Sessions Effect* | *Interaction Effect* |
| *Tot distance moved* | **F (3, 28) = 10.18**<br>**P=0.0003** | **F (2.709, 75.85) = 9.474**<br>**P<0.0001** | **F (9, 84) = 4.947**<br>**P<0.0001** |
| *N° of entrances* | **F (3, 28) = 14.21**<br>**P<0.0001** | **F (2.419, 67.73) = 5.921**<br>**P=0.0025** | **F (9, 84) = 3.969**<br>**P=0.0003** |
| *Latency to first entrance* | **F (3, 28) = 10.46**<br>**P<0.0001** | **F (2.588, 72.48) = 5.856**<br>**P=0.0020** | **F (9, 84) = 2.977**<br>**P=0.0040** |
| *Time spent in target sector* | **F (3, 28) = 7.028**<br>**P=0.0011** | **F (2.797, 78.31) = 7.362**<br>**P=0.0003** | **F (9, 84) = 4.259**<br>**P=0.0001** |

Furthermore, two-way RM ANOVA comparing Rev3 with set-shifting session revealed a significant effect of the groups in the total distance moved, in the n° of entrances, in the latency to enter the target sector, and the time spent in the target sector. Also, a significant effect of the days was found in the four behavioral parameters. Multiple comparisons suggest that *γ*-like 50Hz optogenetic stimulations in the mPFC had an immediate and generally lasting effect in flexibly shifting the RF-to-AF navigation rule compared to the WT and PV-Cre-vHPC groups. However, the vHPC-PV-Cre group showed a better ability to shift the RF-to-AF navigation rule, spending less time in the target sector than the mPFC-Pv-Cre group. Conversely, the mPFC-PV-Cre group showed a better ability to “re-learn” the new navigation rule during the set-shifting days than the vHPC-PV-Cre group.

### Comparisons between optogenetic stimulations and pharmacological control groups (NaCl vs. MK801): a resemblance with the NaCl groups

Results are summarized in Table 4 (Tab. 4). A two-way RM ANOVA on reversal session revealed an effect of the groups in the total distance moved, n° of entrances, in the latency to enter the target sector, and the time spent in the target sector, suggesting that WT-mPFC group resembled MK801 group, and that mPFC-PV-Cre and NaCl behaved similarly. Furtherly, it has been found an effect of the days in the n° of entrances and in the time spent in the target sector suggesting that both PV-Cre groups (mPFC and vHPC) and NaCl group showed an ability to reverse the previous spatial memory throughout the days. A two-way RM ANOVA on set-shifting revealed an effect of the groups in the total distance moved, in the n° of entrances, in the latency to enter the target sector, and the time spent in the target sector. Multiple comparisons suggested that *θ*-like 10Hz optogenetic stimulations in the PV-Cre vHPC group did not produce any overall resemblance to the NaCl group. However, as previously mentioned, PV-Cre vHPC showed similarities with the NaCl group for the time spent in the target sector – a measure of flexible shifting of the RF-to-AF navigation rule. Furtherly, it has been found an effect of the days in the n° of entrances, in the latency to enter the target sector, and in the time spent in the target sector. Multiple comparisons in the time spent in the target sector suggested a general ability to learn the new navigation rule in WT-mPFC and MK801 groups. At the same time, the PV-Cre groups (mPFC and vHPC) and NaCl group were not able to increase the shifting ability throughout the days, furtherly suggesting that both *γ*/*θ*-like optogenetic stimulations induced similar cognitive abilities to the naïve NaCl group.

**Table 4.**
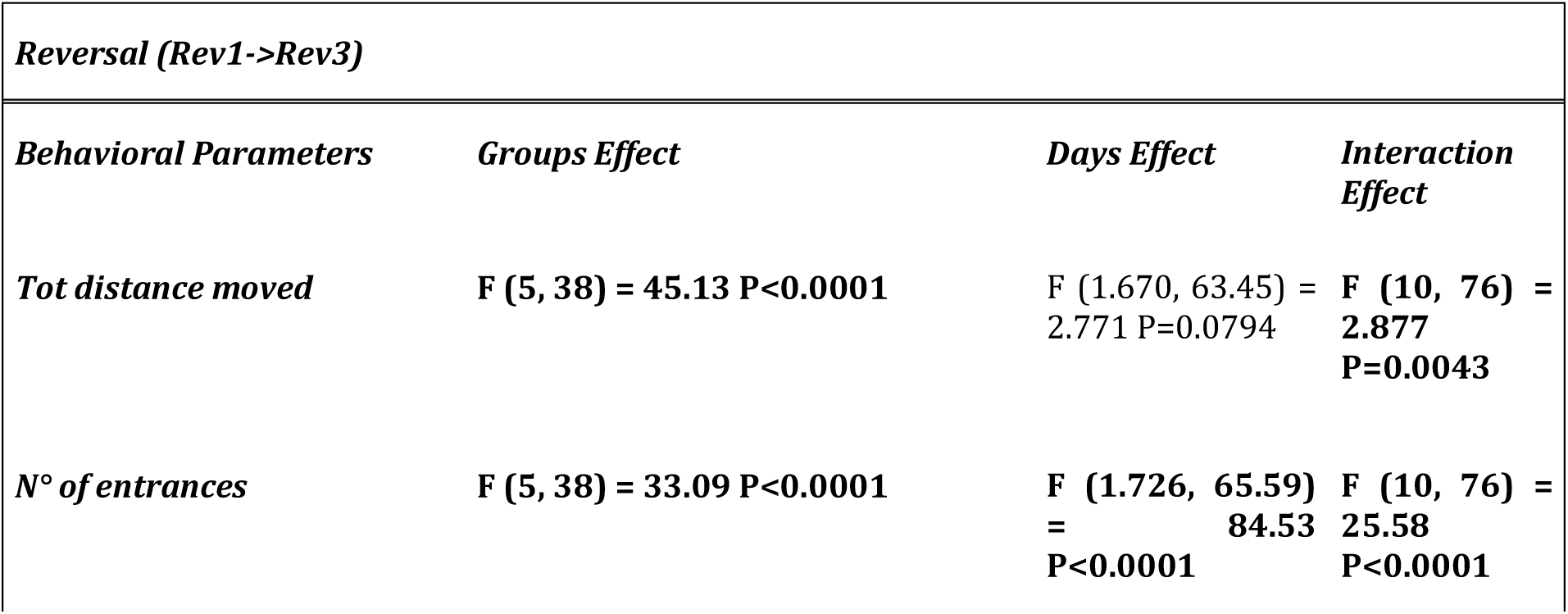
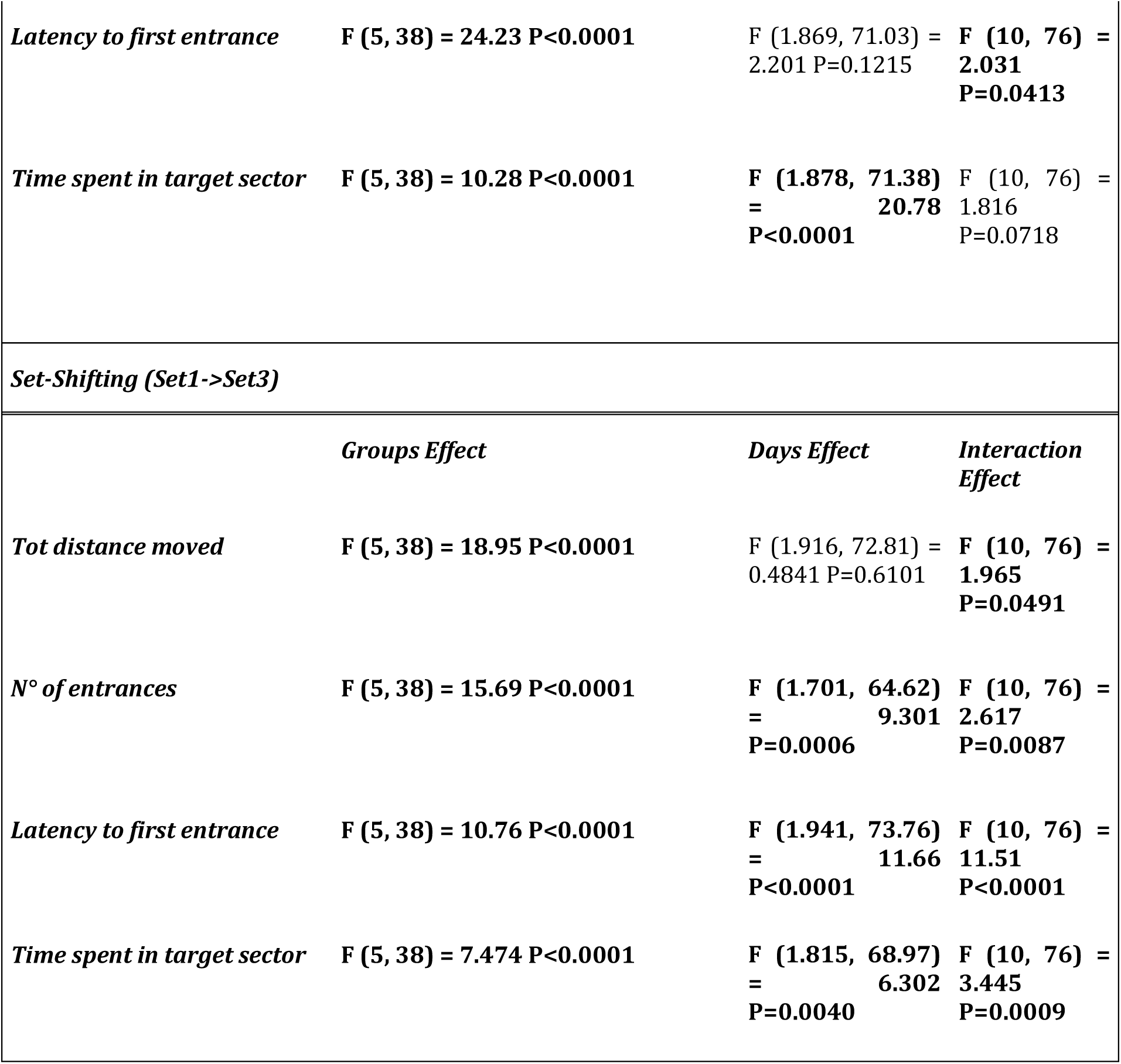
Summary of the two-ways RM ANOVAs of comparisons between optogenetic stimulations and pharmacological control groups during reversal and set-shifting sessions. Significant values are in bold.

### Comparisons with NaCl control groups: ruling out potential effects of MK801 systemic injection

As previously mentioned, all the experimental and the control groups received NaCl instead of MK801 systemic injections and optogenetic stimulations during the reversal and the set-shifting days. Results are summarized in Table 5 (Tab. 5). A two-way RM ANOVA on reversal session revealed an effect of the groups in the total distance moved, in the n° of entrances, in the latency to enter the target sector, and in the time spent in the target sector, thus suggesting that NaCl systemic injections-optogenetic stimulations pairings worsened the inflexibility, especially in the vHPC groups. An effect of the days was found in the latency to enter the target sector and the time spent in the target sector, thus confirming that combinations of NaCl i.p. and optogenetic stimulations of PV+ interneurons were ineffective in restoring reversal spatial memory. Subsequently, a two-way RM ANOVA on set-shifting revealed an effect of the groups in the total distance moved, in the n° of entrances, in the latency to enter the target, and in the time spent in the target sector, thus confirming that MK801-mPFC group performed better than its NaCl group in all the behavioral parameters. These results suggest that the combination of NaCl systemic administration and optogenetic stimulation of PV+ interneurons in mpFC and vHPC exacerbate navigational inflexibility.

**Table 5.** Summary of the two-ways RM ANOVAs of comparisons between MK801 and NaCl groups during PV+ optogenetic stimulations during reversal and set-shifting sessions. Significant values are in bold.

|  |  |  |  |
| --- | --- | --- | --- |
| ***Reversal (Rev1->Rev3)*** |  |  |  |
| ***Behavioral Parameters*** | ***Groups Effect*** | ***Days Effect*** | ***Interaction Effect*** |
| *Tot distance moved* | **F (3, 20) = 11.29 P=0.0002** | F (1.443, 28.86) = 2.373 P=0.1244 | **F (6, 40) = 3.129 P=0.0131** |
| *N° of entrances* | **F (3, 20) = 8.334 P=0.0009** | F (1.807, 36.13) = 0.1474 P=0.8429 | F (6, 40) = 0.1045 P=0.9955 |
| *Latency to first entrance* | **F (3, 20) = 4.929 P=0.0101** | **F (1.616, 32.31) = 8.535 P=0.0020** | F (6, 40) = 0.2884<br>P=0.9390 |
| *Time spent in target sector* | **F (3, 20) = 8.161 P=0.0010** | **F (1.788, 35.76) = 14.71 P<0.0001** | F (6, 40) = 1.702<br>P=0.1458 |
| ***Set-Shifting (Set1->Set3)*** |  |  |  |
|  | ***Groups Effect*** | ***Days Effect*** | ***Interaction Effect*** |
| *Tot distance moved* | **F (3, 20) = 7.177 P=0.0019** | F (1.466, 29.33) = 0.5616 P=0.5239 | F (6, 40) = 1.440<br>P=0.2236 |
| *N° of entrances* | **F (3, 20) = 8.737 P=0.0007** | F (1.614, 32.29) = 0.9561 P=0.3777 | F (6, 40) = 1.142<br>P=0.3562 |
| *Latency to first entrance* | **F (3, 20) = 24.84 P<0.0001** | F (1.698, 33.97) = 2.308 P=0.1220 | **F (6, 40) = 2.440 P=0.0419** |
| *Time spent in target sector* | **F (3, 20) = 12.11 P<0.0001** | F (1.745, 34.90) = 0.8601 P=0.4183 | F (6, 40) = 0.3788<br>P=0.8881 |

### Effectiveness of the viral transfections and confirmation of the local optogenetic stimulations

Firstly, we counted the PV+ cells in the region of interest (ROI, 10mm^2^) as % of the fluorescent area (% area) at rostro-caudal layers (+2.1 to +1.3 AP with respect to bregma, mPFC; −2.7 to −3.5 with respect to bregma, vHPC). A RM one-way ANOVA found a significant effect of optogenetic stimulations in mPFC (Fig 6a), and vHPC (Fig. 6b). Comparisons showed a significant difference between the caudal and rostral slices and the optic fiber position in mPFC and vHPC. Further, we analyzed the percentage of the area (% area) (Fig. 6c) and the integrated density (integrated density) of fluorescence (area x mean gray value) for ChR2/PV+ interneurons and c-Fos protein expressions (Fig. 6d). For the % area, a one-way ANOVA showed an effect of the optogenetic stimulations. Multiple comparisons interestingly showed that mPFC-WT group had a significative increase of c-Fos expression despite the ineffectiveness of PV+ optogenetic stimulations. Also, a significative difference between the expression of PV+/ChR2 and c-Fos protein in the vHPC-PV-Cre group was found, suggesting that despite the optogenetic stimulations, PV+ were not activated by those stimulations (Fig. 6c). A one-way ANOVA for the integrated density substantially replicated the previous results for the % of the area. Multiple comparisons confirmed that despite the optogenetic stimulations, PV+ were not activated (Fig. 6d). Of note, the expression of c-Fos protein in the WT groups of both mPFC and vHPC was similar to that of PV-Cre groups, ruling out potential adverse effects of optogenetic stimulations. Moreover, we analyzed the level of colocalization of ChR2/PV+ interneurons and c-Fos protein around the tip of the optic fiber (ROI, 1mm^2^). A one-way ANOVA analysis showed an extreme significance in both PV-Cre groups, suggesting that optogenetic stimulations indeed activated PV+ interneurons (Fig. 6e). In Figure 6e and 6f, representative areas and cells of mPFC and vHPC, respectively, are shown.

## Discussions

This study sought to determine if optogenetic stimulations of PV+ interneurons at specific frequencies in specific brain regions could restore navigation flexibility using a task that measured reversal spatial memory and navigational set-shifting.

### Comparisons between mPFC and vHPC: the role of optogenetic stimulation

In our first set of experiments, we compared mPFC and vHPC groups (WT and PV-Cre) on acquisition, reversal spatial memory and navigational set-shifting sessions (Fig. 2, Suppl. Mat.). Also, we compared the last day of acquisition (Acq3) with the reversal session days (Acq3 vs. Rev1->Rev3), and the last day of the reversal session with the set-shifting session days (Rev3 vs. Set1->Set3) (Fig. 3). We showed that optogenetic stimulations of PV+ interneurons at specific frequencies in specific brain regions can discretely restore navigational flexibility. Specifically, during reversal days, γ-like (50Hz) stimulations of PV+ in mPFC had a significant effect on the ability to reverse the previous spatial memory as for the total distance moved and the latency to enter the target sector compared to the WT group (Fig. 2e, g; Suppl. Mat.). On the other hand, as for the time spent in the target sector, which is considered a measure of how flexibly mice can reverse the spatial memory of the previous to-be-avoided sector, the γ-like stimulations in mPFC affected the reversing ability compared to the WT group and to the vHPC-PV-Cre group (receveing θ-like optogenetic stimulations of PV+) (Fig 2h; Suppl. Mat.). This last result suggests that putatively, γ-like optogenetic stimulations of PV+ interneurons in mPFC have a more crucial role in reversing spatial memory than θ-like optogenetic stimulations of PV+ in vHPC. Previous studies showed that vHPC is crucial for navigation and flexible reversal of spatial memory [36–37], and that θ oscillation synchronization has a crucial role in these cognitive abilities [38–39]. However, studies reported that somatostatin-positive (SST+) interneurons, rather than PV+ interneurons, gate the vHPC inputs to the mPFC to encode spatial information in the spatial working memory task [40]. In that study, optogenetic inhibition of SST+ and PV+, rather than stimulation, and no pharmacological animal model was used. Moreover, the spatial working memory task consisted of a T-maze with baited arms. Taken together, our results are in line with previous studies and, most importantly, put a link between the role of mPFC-γ/PV+ and vHPC-θ/PV+ oscillation synchronization in flexible reversal memory, that is, a high-order cognitive ability, compromised in SCZ-like conditions.

**Figure 1:**
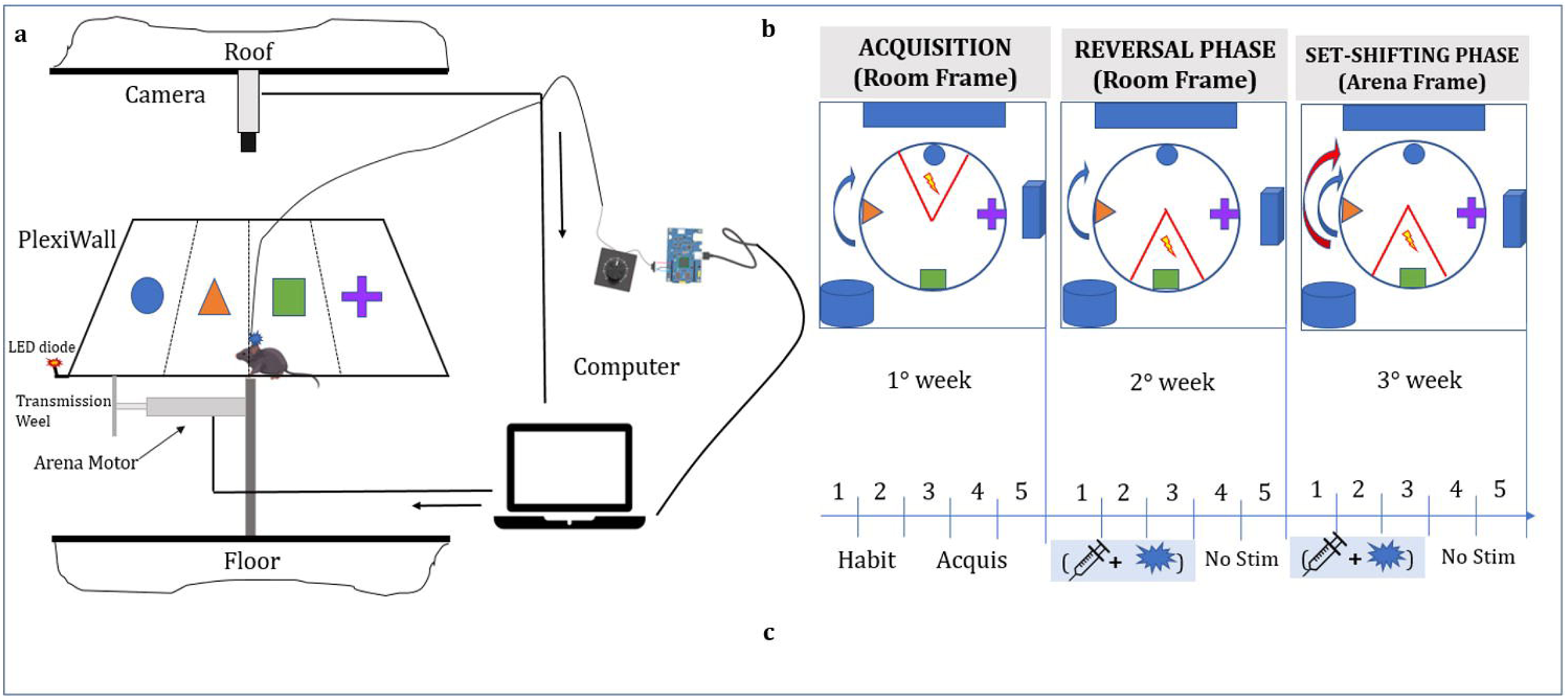
The active place avoidance task. (a) represents a drawing of the place avoidance setup, including the optogenetic stimulation setup. The mouse head optic fiber was connected to a patch-cord, which was cabled to a blue-LED driver, and then to an Arduino board and a laptop. An open-source software (Bonsai.org) controlled LED light pulsing. (b) shows a schematic representation of the three phases (acquisition, reversal, and set-shifting). Note that acquisition and reversal sessions used the room frame (RF) setup rule, while the set-shifting used the arena frame (AF) setup rule. (c) shows the timeline of the procedure. In the first week, mice had 2 days (20 min sessions) to habituate to the arena. Here, only the arena was moving and no footshock was delivered. After, mice had 3 days to acquire the avoidance of the target sector, with rotating arena and footshock (0.3-0.6mA, 500 mmsec/2secs). In the second week, reversal sessions had mice for 3 days with MK801 i.p. and PV+ optogenetic stimulations combinations, to reverse the previous spatial acquisition. Finally, in the third week, set-shifting sessions had mice for 3 days with MK801 i.p. and PV+ optogenetic stimulations combinations to switch the the RF-to-AF rule.

**Figure 2:**
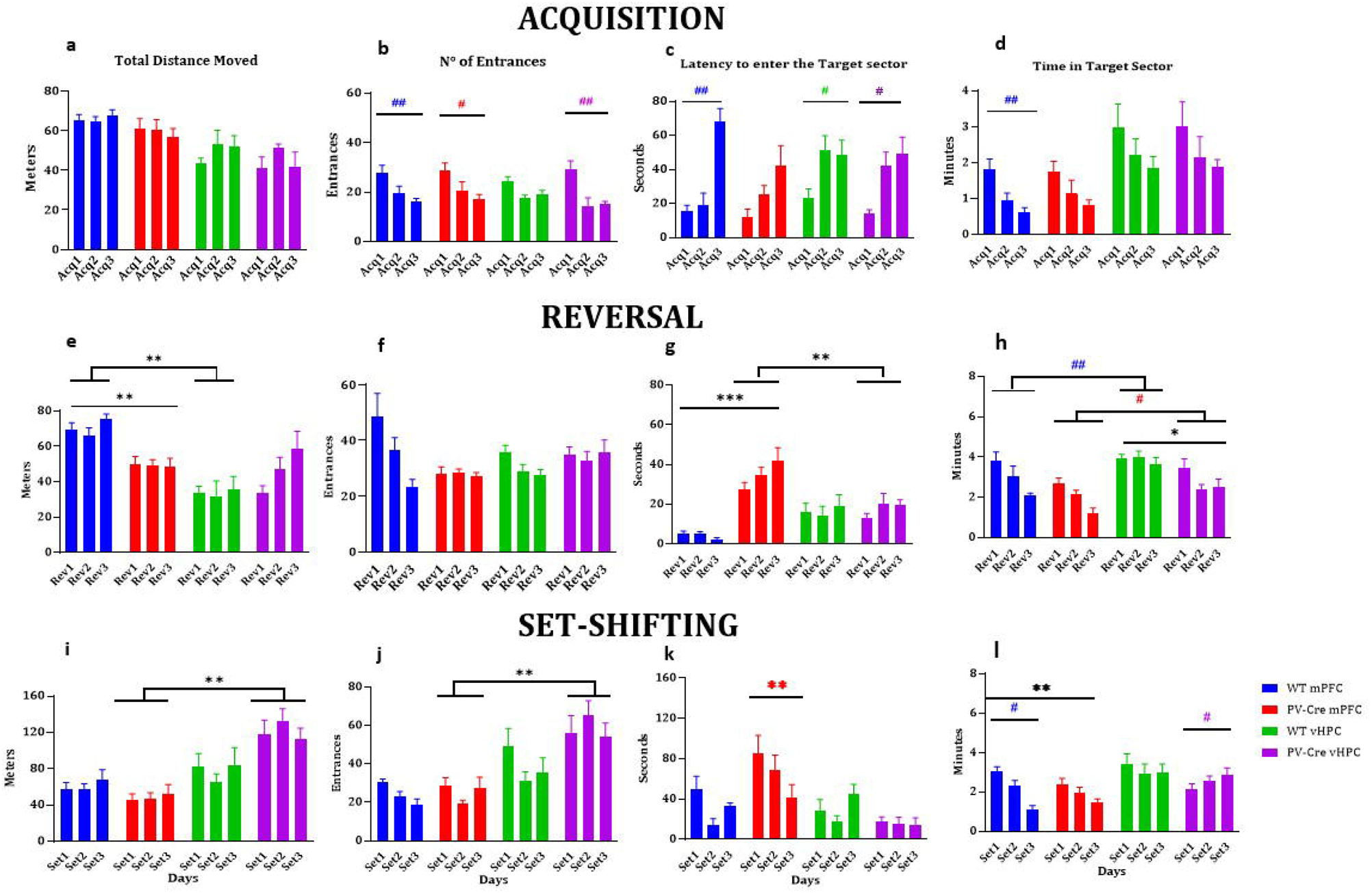
Comparisons between mPFC and vHPC. The first row of panels shows the behavioral results during acquisition sessions for the total distance moved (a), the n° of entrances (b), the latency to enter the target sector (c), and the time spent in the target sector (d). In this session, mice did not receive MK801 i.p. nor optogenetic stimulations. Overall, all groups showed an ability to spatially acquire the position of the target sector during the days (# are colored according to the bars representing the groups) (the sector that must be avoided) (#p value <0.05; ##p value <0.01) (# are colored according to the bars representing the groups, indicating a difference between the days). The second row of panels shows the behavioral results during reversal sessions for the total distance moved (e), the n° of entrances (f), the latency to enter the target sector (g), and the time spent in the target sector (h). In this session, mice received MK801 i.p. and optogenetic stimulations. PV+ interneurons optogenetic stimulations had an effect in the total distance moved (e) and in the latency to enter the target sector (g) between WT and PV-Cre groups of mPFC, and against the WT group of vHPC (e), and between WT and PV-Cre groups of mPFC, and against the PV-Cre group of vHPC (g) (**p value <0.01; ***p value <0.001). As for the time spent in target sector (h), mPFC groups showed a better ability to reverse the previous spatial memory of the target sector during the days compared to their respective groups of vHPC (# are colored according to the bars representing the better groups compared with their respective groups) (*p value <0.05; #p value <0.05; ##p value <0.01). The third row of panels shows the behavioral results during set-shifting sessions for the total distance moved (i), the n° of entrances (j), the latency to enter the target sector (k), and the time spent in the target sector (l). In this session, mice received MK801 i.p. and optogenetic stimulations. PV+ interneurons optogenetic stimulations had an effect in the total distance moved (i) and in the n° of entrances (j) between PV-Cre groups (mPFC vs. vHPC) (**p value <0.01). As for the latency to enter the target sector (k), mPFC-PV-Cre group showed a better ability to avoid entering in the target sector compared to the other groups (* is colored according to the bars representing the better group) (**p value <0.01). As for the time spent in target sector (l), mPFC-PV-Cre group showed a better ability to switch the RF-to-AF navigation rule compared to the WT group (**p value <0.01). Conversely, mPFC-WT group showed a better ability to acquire the new rule (#p value <0.05). Finally, the vHPC-PV-Cre group showed instead the inability to switch the navigation rule between the days (#p value <0.05). Error bars represent ± SEM.

**Figure 3:**
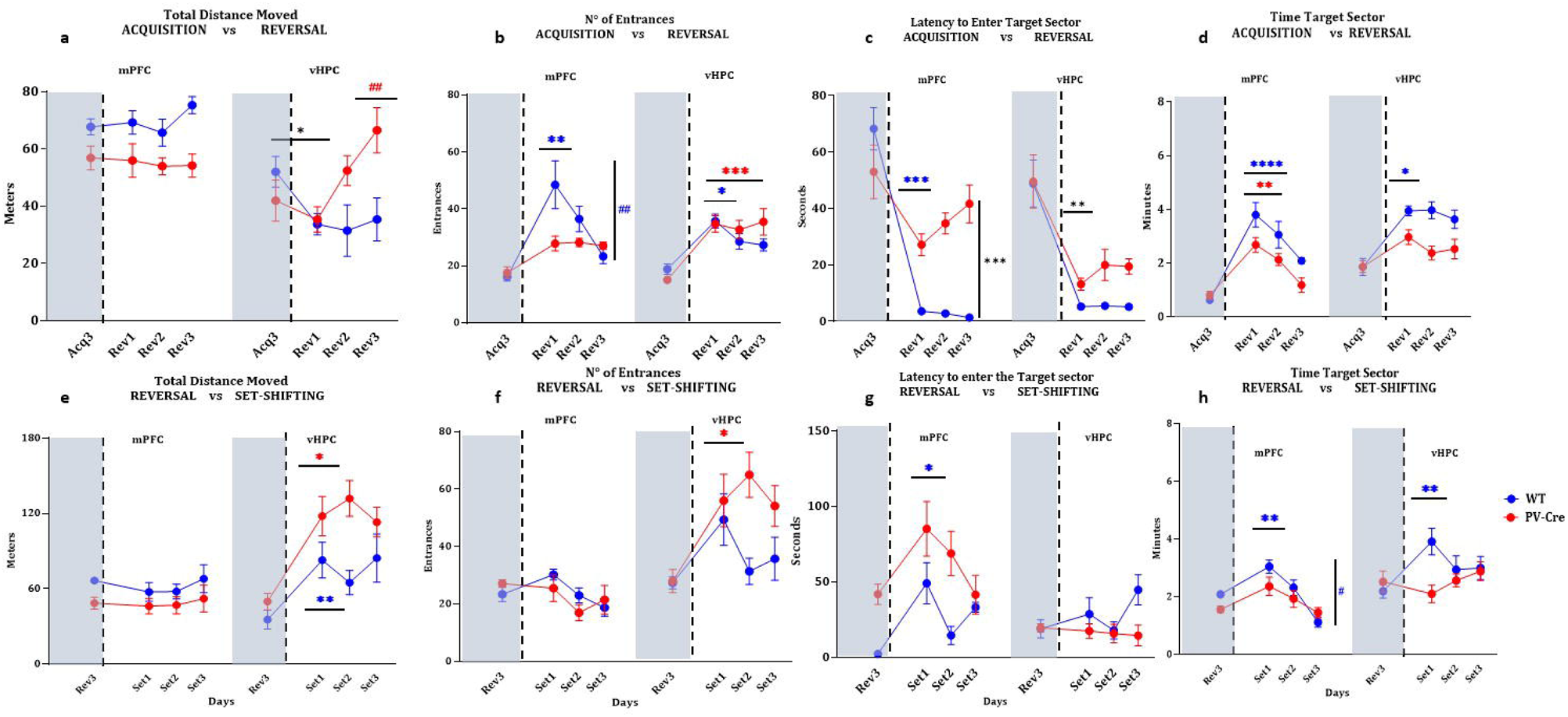
The rescuing role of PV+ optogenetic stimulations. The first row of panels shows comparisons between the last day of acquisition (Acq3) and the reversal days (Rev1, Rev2, Rev3) for the total distance moved (a), the n° of entrances (b), the latency to enter the target sector (c), and the time spent in the target sector (d). Both WT groups showed an effect of the MK801 i.p. administration for all the behavioral parameters measured, except for the mPFC-WT group in the total distance moved (a) (*p value <0.05; **p value <0.01; ***p value <0.001; ****p value <0.0001; colored asterisks are in accordance with the colored lines representing the groups, while black asterisks correspond to comparisons between groups). Moreover, the mPFC-WT group showed an ability to learn to avoid entering in the newly forbidden sector (b) (##p value <0.01; # is colored according to the correspondent line representing a group of mice). On the other hand, the PV-Cre groups discretely showed an effect of PV+ optogenetic stimulations. The mPFC-PV-Cre group showed an effect of optogenetic stimulation for all behavioral parameter measures, except for the time spent in target sector (d) (**p value <0.01; ***p value <0.001; ****p value <0.0001; colored asterisks are in accordance with the colored lines representing the groups). Conversely, the vHPC-PV-Cre group showed an effect of the optogenetic stimulations only for the time spent in the target sector (d) (*p value <0.05). The second row of panels shows comparisons between the last day of reversal (Rev3) and the set-shifting days (Set1, Set2, Set3) for the total distance moved (e), the n° of entrances (f), the latency to enter the target sector (g), and the time spent in the target sector (h). The mPFC-WT group showed an effect of the MK801 i.p. administration over the ability to switch the navigation rule for the latency to enter the target sector (g) and the time spent in target sector (h), and in the total distance moved (e) (*p value <0.05; **p value <0.01). However, as for the time spent in target sector, the mPFC-WT showed an ability to learn the rule switching throughout the set-shifting days (#p value <0.05). Finally, the vHPC-WT group showed an effect of the MK801 administration for the total distance moved (e) and the time spent in target sector (h), while the vHPC-PV-Cre group showed an effect of the optogenetic stimulation for the latency to enter in target sector (g) and for the time spent in target sector (h) (**p value <0.01; ***p value <0.001; colored asterisks are in accordance with the colored lines representing the groups). Error bars represent ± SEM.

During set-shifting days, again, γ-like stimulations in mPFC had a significant effect on the ability to switch the RF-to-AF navigation rule compared to θ-like stimulations in vHPC in the total distance moved and the n° of entrances (Fig. 2i, j; Suppl. Mat). As for the latency to enter the target sector, γ-like PV+ optogenetic stimulations in the mPFC-PV-Cre group restored the ability to step into the target sector compared to all the other groups (Fig. 2k; Suppl. Mat.). Furthermore, as for the time spent in the target sector, differences between mPFC groups (WT vs. PV-Cre) were reported (Fig. 2l; Suppl. Mat.). On the other hand, a significative difference was reported between mPFC and vHPC-PV-Cre groups on day 3 of the set-shifting, showing that mPFC-PV-Cre group had a better ability to learn the switch of the navigation rule, thus suggesting that, putatively, γ-like stimulations in mPFC had a significant role in rule-switching than θ-like stimulations in vHPC (Fig. 2l; Suppl. Mat.). Previous studies demonstrated that mPFC is crucial in flexible rule-switching, which is cognitive flexibility based on problem-solving and decision-making [7, 41], and that γ oscillation synchronization in mPFC has a significant role in cognitive functions [21, 23, 42], including behavioral flexibility [2]. Altogether, our results align with previous studies, suggesting a specific role for specific wave oscillations in specific brain regions for specific cognitive abilities, which is pivotal in the mPFC-vHPC compromised crosstalk in SCZ-like conditions.

Comparing Acq3 vs. Rev1->Rev3, we showed that γ-like stimulations in mPFC and θ-like stimulations in vHPC, affected differently the ability to restore the spatial memory, as measured by the n° of entrances, the latency, and the time spent in the target sector (Fig. 3a, b, c). Considering the time spent in the target sector during reversal sessions as a measure of flexibility (the more time spent in the newly to-be-avoided sector, the more inflexibility), only vHPC-PV-Cre group showed an optogenetically-induced flexibility (Fig. 3d). Furthermore, comparisons in Rev3 vs. Set1->Set3 showed that γ-like stimulations in mPFC significantly restored the total distance moved and the n° of entrances (Fig. 3e, f). On the other hand, the θ-like stimulations in vHPC influenced the latency to enter the target sector and the time spent in the target sector (Fig. 3g, h). The Rev3 vs. Set1->Set3 comparisons for the time spent in the target sector showed that both stimulations (γ-like in mPFC and θ-like in vHPC) restored navigation set-shifting compared to their own WT groups. Altogether, these results confirm the previous behavioral outcomes, thus suggesting a pivotal role of optogenetic stimulations of PV+ interneurons at specific frequencies in specific brain regions, which are considered crucial for executive functions and spatial navigation [2, 21, 36–37, 42].

### Comparisons with non-optogenetically stimulated groups (NaCl vs. MK801): a resemblance with the NaCl group

In the second set of experiments, we sought to determine whether the “optogenetic rescue” of navigational flexibility was comparable to the ones of healthy subjects. To this aim, we compared the “optogenetic rescue” of reversal spatial memory and navigation set-shifting, previously altered by MK801systemic administration, with groups of mice only receiving MK801 i.p administration (compared with WT groups), or NaCl i.p. administration (compared with PV-Cre groups). Comparisons of the reversal days showed that the PV-Cre groups (mPFC and vHPC) had similar abilities to the NaCl group in the total distance moved and the n° of entrances (Fig 4a, b). Notably, the mPFC-PV-Cre group resembled the NaCl group regarding the latency to enter the target sector and the time spent in it (Fig. 4c, d). Seemingly, the mPFC-WT group resembled the MK801 group in the inability to restore reversal spatial memory for the n° of entrances, the latency to enter the target sector, and the time spent in the target sector (Fig. 4b, c, d). Conversely, comparing the set-shifting days, the ability to restore navigation flexibility by specific frequencies of optogenetic stimulations was more visible in mPFC-PV-Cre group than the vHPC-PV-Cre compared with the NaCl group (Fig 4e, f, g). However, it is essential to note that the navigation flexibility was better restored in both the PV-Cre groups (mPFC and vHPC) than in the NaCl group, although not significantly (Fig. 4h). Nevertheless, taken together, these results demonstrate that specific optogenetic stimulations in specific brain regions might have a therapeutical role in restoring navigation flexibility. Previous studies have shown the optogenetic ability to restore behavioral flexibility [43–44], and particularly, the role of optogenetic stimulations of PV+ interneurons [2].

**Figure 4:**
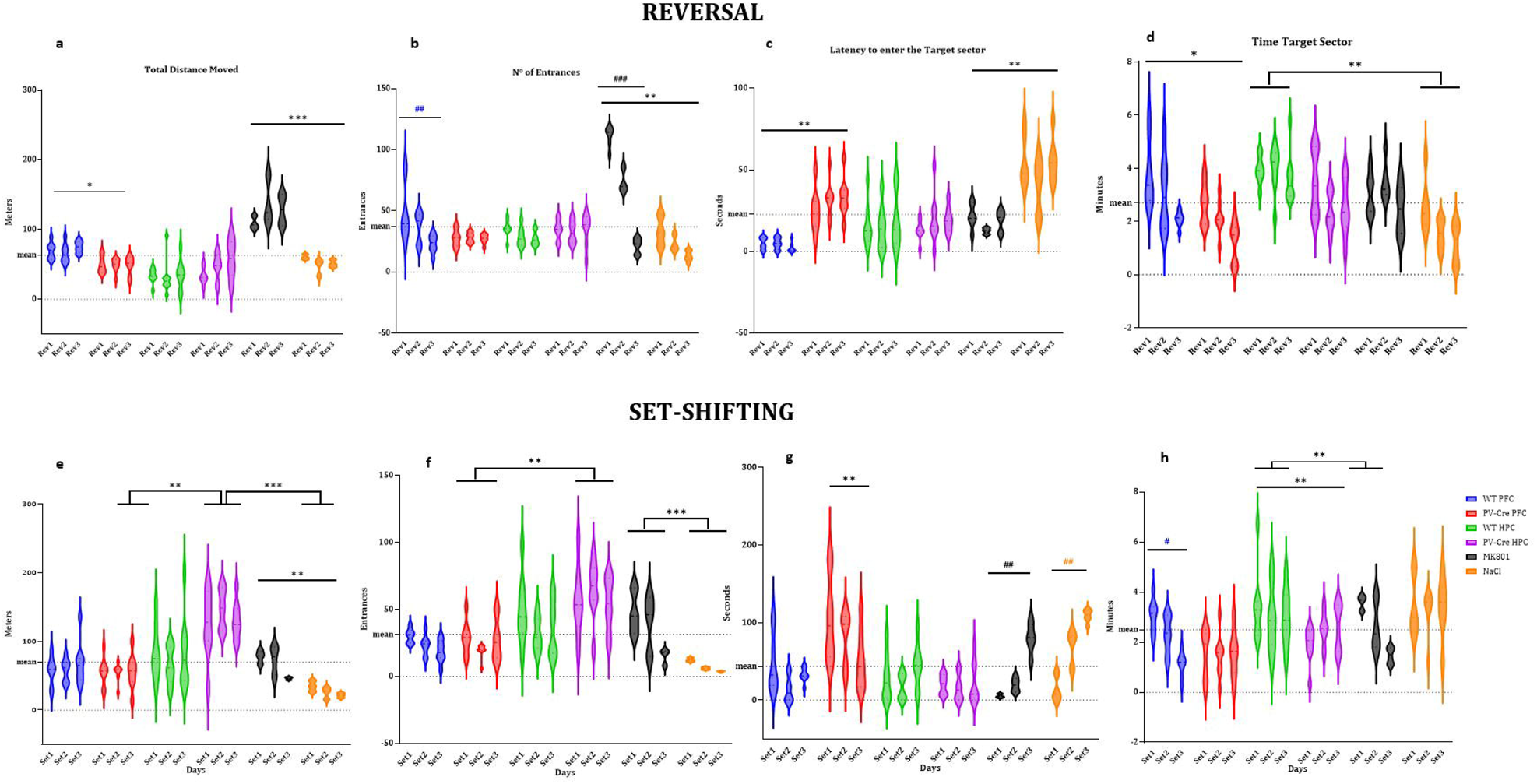
Comparisons between optogenetic stimulations and pharmacological control groups. With these comparisons, we sought to determine whether PV+ optogenetic stimulations in mPFC and vHPC restored the navigational flexibility to the level of NaCl control groups. The first row of panels shows the behavioral results during reversal sessions for the total distance moved (a), the n° of entrances (b), the latency to enter the target sector (c), and the time spent in the target sector (d). Mice receiving only MK801 i.p. (0.0.8mg/Kg) showed a pattern of navigational inflexibility similar to the mPFC-WT group, but not the vHPC-WT group for the total distance moved (a), the n° of entrances (b), and the latency to enter the target sector (c). As for the time spent in the target sector (d), the vHPC-WT showed a significative inability to reverse the previously acquired to-be-avoided sector compared to the NaCl group. Overall, only the mPFC-PV-Cre group showed a behavioral pattern of restored navigational flexibility similar to the NaCl group (*p value <0.05; **p value <0.01; ***p value <0.001; ##p value <0.01; ###p value <0.001; # are colored according to the violins representing the groups, indicating a difference between the days). The second row of panels show the behavioral results during set-shifting sessions for the total distance moved (e), the n° of entrances (f), the latency to enter the target sector (g), and the time spent in the target sector (h). In this session, the comparisons between the optogenetically stimulated groups confirmed that mPFC-PV-Cre group rescued the total distance moved (e), the n° of entrances (f), and the latency to enter the target sector (g) compared to vHPC-PV-Cre group, while vHPC-PV-Cre showed a better ability to switch the navigational rule than the mPFC-PV-Cre group (h). As for the comparisons with the pharmacological groups (MK801 and NaCl), WT groups showed a major resemblance with the MK801 group than the PV-Cre groups with the NaCl group **p value <0.01; ***p value <0.001; ##p value <0.01; ###p value <0.001; # are colored according to the violins representing the groups, indicating a difference between the days). Dotted lines represent the 0 and the grand mean of all groups. Error bars represent ± SEM.

### Comparisons with optogenetically-stimulated NaCl control groups: ruling out potential effects of MK801 systemic injection

In the third set of experiments, we wanted to determine whether MK801 i.p. administrations *per se* were affecting the behavioral outcomes, and whether optogenetic stimulations were properly effective in counteracting the MK801 administration. Performing a new set of experiments using NaCl i.p. administration in new mPFC and vHPC groups (WT and PV-Cre), we compared them with the previous groups (MK801 i.p.+optogenetic stimulations) in reversal and set-shifting sessions. The results are shown in Fig. 5 demonstrating the validity of the acute MK801 model of SCZ-like inflexibility and showing that specific optogenetic stimulations of PV+ interneurons in specific brain areas can restore the previous MK801-induced inflexibility. Furthermore, this set of comparisons reveals that the combinations of NaCl i.p. administrations and PV+ optogenetic stimulations did not ameliorate the restore of navigational flexibility. Rather, the NaCl i.p.-PV+ optogenetic stimulations combination did exacerbate navigational inflexibility (Fig. 5e, f, g, h). A possible explanation for these results is that being MK801 hypofunctioning NMDARs expressed on PV+ interneurons [45–46], NaCl i.p. administration did not produce any NMDARs hypofunctioning effect. In addition, one of the most accepted hypotheses is the E/I altered balance in SCZ and other diseases affecting cognitive abilities [19, 47–49]. In the “classical” condition, when modeling acute SCZ-like cognitive deficits, MK801 is systemically administered acutely, inducing the E/I altered ratio and the desynchronization of γ and θ bands oscillations [50–52]. Optogenetic stimulations of PV+ interneurons have the role of putatively “rebalancing” the E/I ratio [2, 44], thus potentially restoring behavioral flexibility. Instead, when NaCl is systemically administered and coupled with optogenetic stimulations of PV+ interneurons, putatively, this induces an E/I altered ratio and, therefore, potential desynchronization of γ and θ bands oscillations in the brain areas involved during behavioral flexibility.

**Figure 5:**
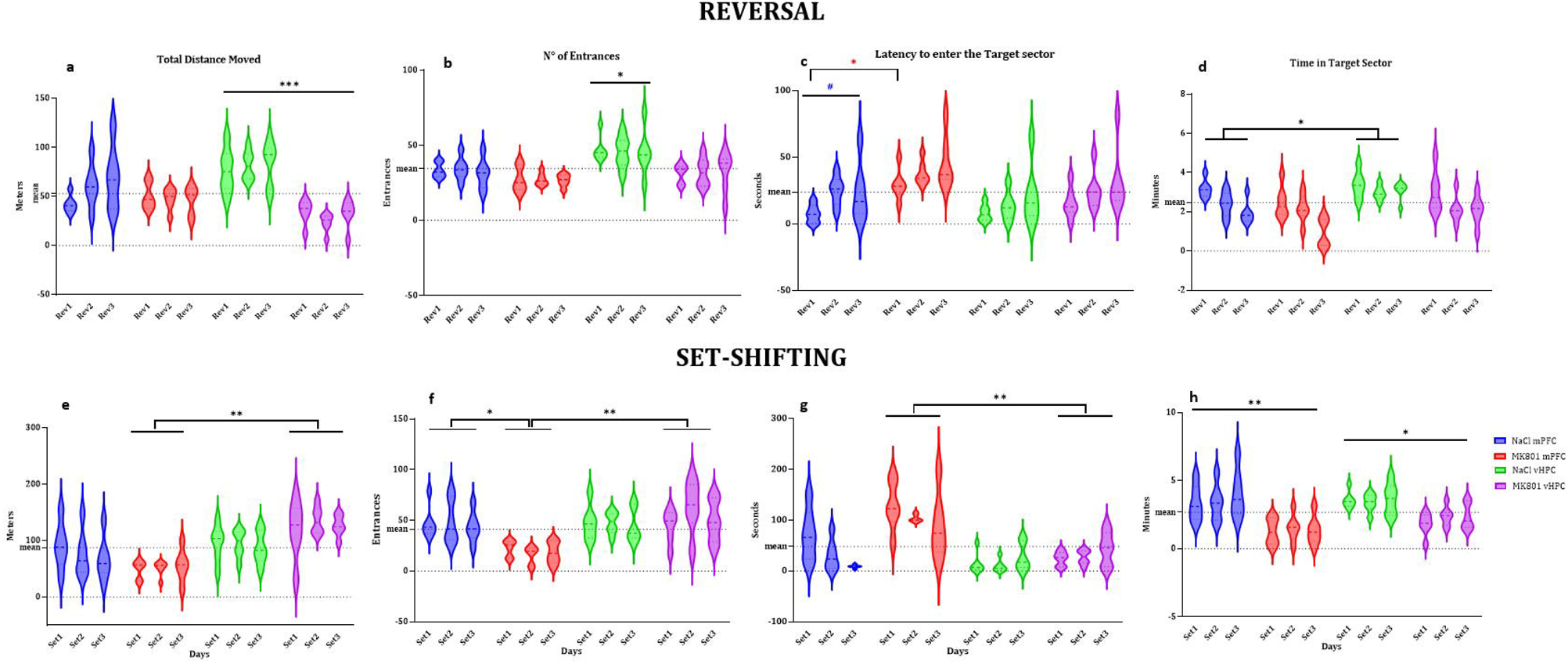
MK801 vs. NaCl comparisons during PV+ optogenetic stimulations. With these comparisons, we sought to determine whether NaCl i.p. administration coupled with PV+ optogenetic stimulations influenced navigational flexibility compared with the “classic” groups having MK801 i.p. administration coupled with PV+ optogenetic stimulations. The first row of panels shows the behavioral results during reversal session for the total distance moved (a), the n° of entrances (b), the latency to enter the target sector (c), and the time spent in the target sector (d). Overall, the NaCl-vHPC group of mice showed a significative decreased ability to reverse the previous spatial localization of the target sector compared to the other groups (*p value <0.05; ***p value <0.001). Only for the latency to enter the target sector (c), the NaCl-mPFC group showed a decreased ability to reverse the location of the previously forbidden sector only in the first day compared to the MK801-mPFC group (*p value <0.05). However, the NaCl-mPFC group also showed an ability to acquire the new location during the days (#p value <0.05; # are colored according to the violins representing the groups, indicating a difference between the days). The second row of panels show the behavioral results during sey-shifting session for the total distance moved (e), the n° of entrances (f), the latency to enter the target sector (g), and the time spent in the target sector (h). Overall, the results confirmed the previous findings regarding the comparisons between mPFC and vHPC groups with MK801-PV+ opto-stimulations for the total distance moved (e), the n° of entrances (f), and the latency to enter the target sector (g). However, as for the n° of entrances (f) and for the time spent in the target sector (h), mice of the NaCl groups (mPFC and vHPC) showed a significative decreased ability to switch the navigation rule (*p value <0.05; ***p value <0.001). Dotted lines represent the 0 and the grand mean of all groups. Error bars represent ± SEM.

### Effectiveness of the viral transfections and confirmation of the local optogenetic stimulations

The image analysis in this study aimed to confirm the effectiveness of the local viral transfections of ChR2 on PV+ interneurons in both mPFC and vHPC and to determine whether optogenetic stimulations induced behavioral flexibility during the reversal and the set-shifting sessions by colocalizing the expression of the c-Fos protein on the PV+/ChR2 interneurons. Counting the % of the area fluorescence in brain slices adjacent to the optic fibers of mPFC and vHPC groups of mice (WT and PV-Cre), we found a significantly higher number of transduced cells around the the tip of the fiber (Fig 6a, 6b), confirming the quality of the surgical procedures of AAV transduction and fiber placement (Fig. 6f, 6g). Moreover, the level of colocalization of c-Fos onto PV+/ChR2 interneurons (Fig. 6c, d, e), suggesting that optogenetic stimulations of ChR2 in vHPC did not sufficiently activate PV+ interneurons. Of note, the expression of c-Fos protein in the WT groups of both mPFC and vHPC was similar to that of PV-Cre groups, ruling out potential adverse effects of optogenetic stimulations. Finally, these results confirm that optogenetic stimulations activated PV+ interneurons. Therefore, this suggests that our optogenetic manipulations affected our measured behavioral parameters, overall rescuing the navigational flexibility.

**Figure 6:**
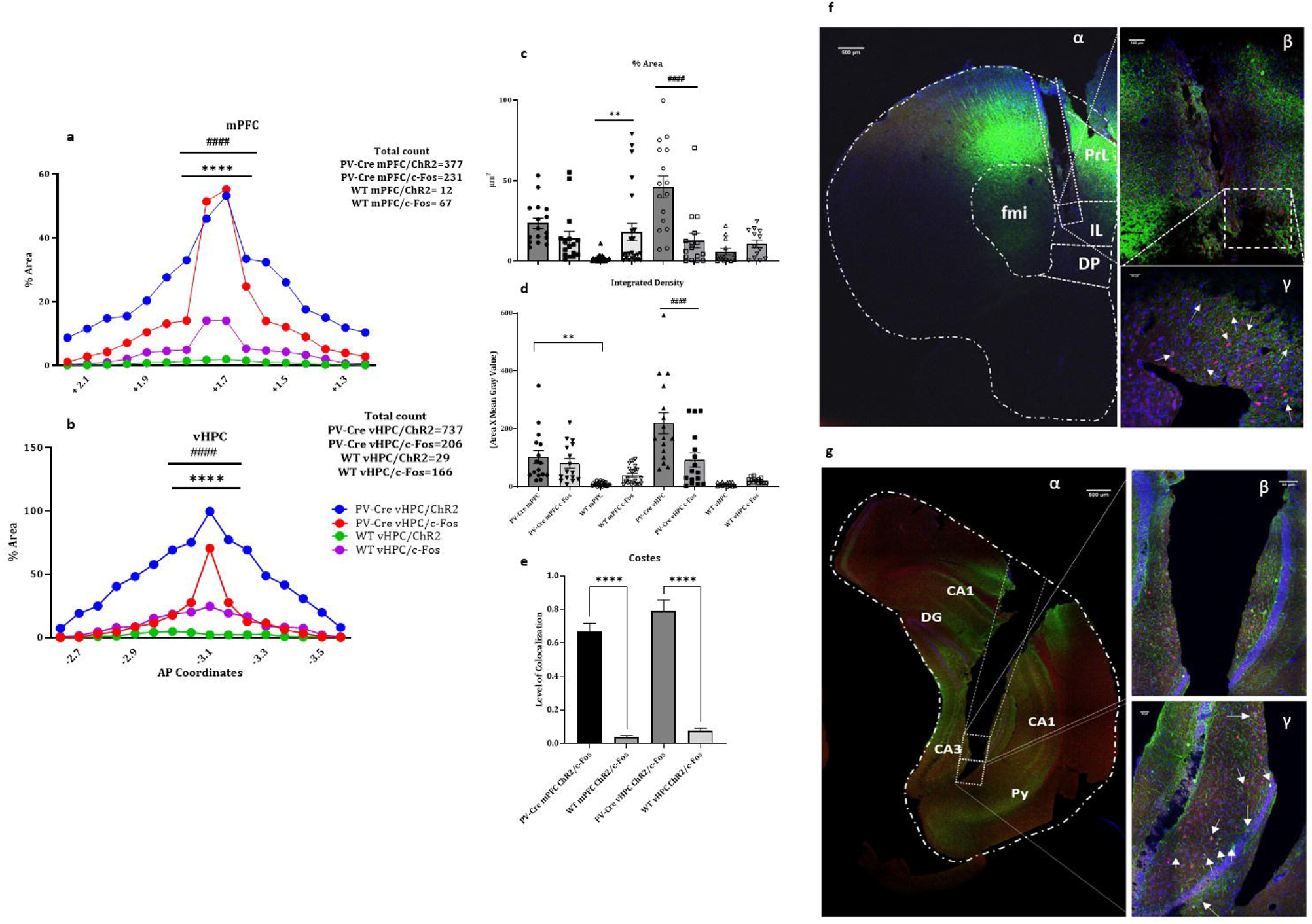
Effectiveness of the optogenetic stimulations. Panels on the left show the results of the % area of fluorescence in the 5 adjacent sections of the mPFC (a) and vHPC (b). The adjacent sections in mPFC cover 0.8 mm (+1.3/+2.1 mm AP to bregma, with the fiber implant at +1.7 mm). The adjacent sections in vHPC cover 0.8 mm (−2.7/-3.5 mm AP to bregma, with the fiber implant at −3.1 mm). Significant effect of optogenetic stimulations in mPFC (a) and vHPC (b) were found with respect to WT groups (****p value<0.0001). Moreover, significant levels of fluorescence were observed in the optic fiber position compared to the other more anterior/posterior slices (####p value<0.0001). Manual counting of the cells is reported in each of the panel. Panels in the center show the % of the area (c) and the integrated density (d) for ChR2/PV+ interneurons and c-Fos protein expressions. Multiple comparisons interestingly showed that mPFC-WT group had a significative increase of c-Fos expression despite the ineffectiveness of PV+ optogenetic stimulations (**p value <0.01), and that a significative difference between the expression of PV+/ChR2 and c-Fos protein in the vHPC-PV-Cre group (####p value<0.0001). Further, lower panel € shows the level of colocalization of ChR2/PV+ interneurons and c-Fos protein around the tip of the optic fiber (ROI, 1mm2). An extreme significance in both PV-Cre groups suggests that optogenetic stimulations indeed activated PV+ interneurons (****p value<0.0001). Panels on the right show representative examples of optic fiber positions in mPFC (f) and vHPC (g) and representative example of colocalization of PV+/ChR2 (in green) and c-Fos protein (in red). Scale bars (fα 500μm, fβ 100μm, fγ 25μm, gα 500μm, gβ 50μm, gγ 25μm).

To conclude, in this study, we aimed to investigate the “rescuing” role of PV+ interneurons optogenetic stimulations in mPFC and vHPC of mice during a task where two high-order executive functions, such as reversal spatial memory and navigation set-shifting were acutely impaired. To our knowledge, this study shows for the first time that a high-order executive function in the form of navigational flexibility is linked to PV+ interneurons activity in specific brain areas such as mPFC and vHPC. Moreover, this study put forward the possibility that optogenetic stimulations of PV+ interneurons in specific brain areas might be putatively linked to specific oscillation waves such as γ (30-80Hz) and θ (8-12Hz), known to be related to cognitive performances [2, 23].

Finally, this study shed light on the complex mPFC-vHPC crosstalk during complex cognitive tasks and the potential new pharmacological target of selective GABAergic agonisms in psychosis treatments. Moreover, this study introduces the question whether disruption of executive functions involves specific loci of brain dysfunction (domain-specific hypothesis) or a systems-level dysfunction (domain-general hypothesis), emphasizing the hierarchical nature of cognitive domains and the need for targeted assessments to differentiate between various psychiatric conditions.

## Supporting information

Suppl. Mat.

## Conflict of Interest Statement

The authors declare no conflict of interest.

## Acknowledgments

This study was supported by Czech Health Research Council grant AZV NU22-04-00526. Image acquisition was supported by IPHYS BIF – MEYS CR (Large RI Project LM2023050 Czech-BioImaging) and ERDF (Project No. CZ.02.1.01/0.0/0.0/18_046/0016045).

## Notes

### Competing Interest Statement

The authors have declared no competing interest.

