## Supplementary material for "PV+ optogenetic stimulations at specific frequencies in specific brain regions can restore navigational flexibility in an acute MK801 mouse model of schizophrenia": Suppl. Mat.

### Supplementary Materials

#### Materials and Methods

***Animals.*** All experiments and animal treatments complied with the Animal Protection Code of the Czech Republic and the European Community Council directive (2010/63/EC). Pilot data showed that our female mice presented high variability in performing the task independently from the estrous cycle's period. Therefore, only male mice (12-15 weeks old) were used. For the optogenetic experiments, the strain Pvalb-T2A-Cre-D mice expressing Cre recombinase directed to PV-expressing cells was used (https://www.jax.org/strain/012358) (PV-Cre). As control, we used wild-type (WT) C57BL/6J mice, commonly used as a general-purpose strain (https://www.jax.org/strain/000664). All the mice for the experiments (N=60) were bred and housed with food and water available ad libitum and maintained on a 12/12 hr light/dark cycle. The experiments were performed during the light phase of the day.

***Drugs.*** Drugs. (+) MK801 (dizocilpine hydrogen maleate; Sigma-Aldrich, CR) was dissolved in 0.9% NaCl (0.08 mg/ml) and injected intraperitoneally at 0.08 mg/kg. The specific dose was chosen to induce cognitive disabilities but not alter locomotion based on pilot data and previous reports [26]. Furthermore, previous studies showed that MK801 dose-dependently affected γ [27] and θ [28] bands, which served as a ground rule in our study to desynchronize and resynchronize γ/θ bands, affecting and restoring cognitive flexibility. Nonetheless, we performed control experiments where NaCl i.p. was injected in PV-Cre mice before optogenetic stimulations to rule out any unspecific effects of the systemic injections. Fresh solutions were prepared on the injection day.

The Cre-dependent viral vector pAAV-Ef1a-DIO -hChR2(E123T/T159C)-EYFP - serotype AAV9 - (ChR2) was a gift from Karl Deisseroth (Addgene plasmid # 35509; http://n2t.net/addgene:35509; RRID: Addgene_35509) [29].

***Active Place Avoidance Test.*** Navigational flexibility was assessed on a rotating arena (Stuchlik et al., 2013) (Fig. 1a). A tracking system recorded the location of the mice and delivered mild shocks (0.3–0.6 mA) if they entered a forbidden 60° sector. Navigation on the arena was guided by various extra-maze cues (e.g., table, cabinet, door) (Room Frame, RF) or distinct cue cards on the arena wall that were different in color and shape (Arena Frame, AF). Arena rotation enabled dissociation between RF and AF frames. All the protocol phases are represented in Fig 1b. In the acquisition session (Acquisition), the shocked sector was static and arbitrarily set to the "north". Thus, the room cues were in the relevant frame (RF) of navigation, and the animals needed to move against the arena rotation to avoid the sector. In the reversal learning session (Reversal), the position of the sector was reversed to the "south", still defined by the cues in the room (RF). The reference frames changed in the navigational flexibility sessions (Set-Shifting), and the sector was set to rotate with the arena (AF). The animals had to start ignoring the irrelevant room cues and pay attention to the arena cue cards, one of which represented the position of the sector. In Fig. 1c, a timeline of the behavioral protocol is represented.

***Surgeries and viral injections.*** All mice weighed ~30 g at the time of the surgery. Unilateral viral infections and optic fiber implants were pseudo-randomly balanced between all subjects' right and left hemispheres. According to the atlas, [30] mPFC coordinates were as follows: AP = +1.8; ML (in 12° angle) = ± 1; DV = -1.8, -2.2 (AAV infections), -2 (optic fiber); and vHPC coordinates were as follows: AP = -3.1; ML = ± 3; DV = -3, -4 (AAV infections), -3.5 (optic fibers). Studies reported that unilateral lesions of mPFC and vHPC are sufficient to affect cognitive abilities [31-32]. In rodents and non-human primates, cognitive functions are not lateralized at the same level as humans. Therefore, cognitive abilities can be supported by communication between intact brain structures in one hemisphere, and unilateral optogenetic stimulations might be sufficient to engage in high-order cognitive tasks.

***Optogenetic stimulation.*** During the reversal and the set-shifting sessions, the mice had the optic fibers plugged into a patch cord delivering a 470 nm light (irradiance, 5-7mW/mm2) at specific frequencies (PFC: 50Hz, 5 ms; vHPC: 10Hz, 10 ms). The stimulation pattern was as follows: 5 min ON, 5 min OFF, 5 min ON, 5 min OFF. We used this pattern of stimulation mimicking "tonic" GABAergic inhibition. Differently from "phasic" GABA inhibition, which transiently desensitizes GABA conductance in cognition [33, for review], "tonic" inhibition is mediated by high affinity-GABA extrasynaptic receptors resulting in a persistent GABAergic conductance, increasing γ/θ band power spectrum for longer than a minute [34].

***Experiments.***

*Experiment 1:* Three weeks after post-surgeries recovery, the four groups of mice (mPFC-WT [n=8], mPFC-PV-Cre [n=8], vHPC-WT [n=8], vHPC-PV-Cre [n=8]) started the behavioral procedures. During the first week of acquisition, the mice were plugged to the optogenetic system, except for the light delivery. During reversal and set-shifting sessions, 30 min prior starting the procedure, mice received MK801 i.p. injections (0.08 mg/Kg). After 30 min, the mice were gently plugged to the optogenetic cable and put in the arena. The optogenetic stimulations were delivered according to the schedule described above. In the last day of set-shifting, the mice were left in the arena for 45 min after finishing the task to allow the potential c-Fos expression. Subsequently, the mice were gently unplugged and rapidly perfused transcardially, and the brains were removed and stored for further IHC validations.

*Experiment 2:* With the aim of comparing the rescuing ability of optogenetic stimulations to control groups, we performed the same behavioral experiment using two groups of mice that only received MK801 (n=8) or NaCl (n=8) i.p. injections during the reversal and set-shifting sessions. During reversal and set-shifting sessions, 30 min prior starting the procedure, mice received MK801 i.p. injections (0.08 mg/Kg). After 30 min, the mice were gently put in the arena.

*Experiment 3:* This experiment resembled the Experiment 1 except that the four groups of mice (mPFC-WT [n=8], mPFC-PV-Cre [n=8], vHPC-WT [n=8], vHPC-PV-Cre [n=8]) received NaCl i.p. injections instead of MK801.

***Immunohistochemical validation.*** Fluorescence immunohistochemical assays were performed to validate c-Fos protein expression on those PV+ interneurons transduced with ChR2. Coronal sections (50µm thickness) of mPFC and vHPC areas were cut using a cryostat (Leica CM1950). Brain sections of both PV-Cre and WT groups were collected and cryopreserved. Fig. 5a and 5b show a representative example of the mPFC and vHPC regions sectioning, with the ChR2 expression around the fiber trace (Fig. 6a, 6b). The collected sections were washed in PBS, permeabilized with PBS and Triton-X 100 1.2% (20 mins, RT), and blocked with PBS and normal bovine serum (NBS) 5% (60 mins, RT). Sections were then incubated in rabbit polyclonal antibodies against c-Fos (Abcam, ab190289, 1:500) and anti-green fluorescent protein (GFP) chicken antibody (ChR2/PV+) (Abcam ab13970, 1:500) in a PBS solution containing 2% NBS and 0.2% Triton X-100 (4°C, overnight). The sections were then washed and incubated in donkey anti-rabbit Alexa Fluor 594 (Invitrogen A32754, 1:500) and anti-chicken Alexa Fluor 488 (Invitrogen A11039, 1:500) secondary antibodies (2 hrs, RT). Brain sections were washed in PBS, mounted onto gelatin-coated slides, and cover-slipped with Vectashield antifade mounting medium containing DAPI (Vector Laboratories, #H1200). Furthermore, images were acquired using a Leica SP8 confocal microscope and analyzed using ImageJ and Fiji plugins. Concerning the quantification of the PV+/ChR2 colocalization, the superposition of fluorescence images is the most prevalent method for evaluating colocalization. The intensity of a pixel in one channel is evaluated against the corresponding pixel in the second channel, generally producing a correlation coefficient. Our colocalization analysis showed a high rate of colocalization among PV+/ChR2 interneurons with c-Fos protein, suggesting that the PV+ were successfully activated by ChR2 when reached by light.

***Data analysis.*** Data were analyzed through ANOVA with repeated measures (RM). Data in figures are represented as mean ± SEM unless otherwise stated. Statistical analysis was performed using GraphPad Prism 9.1.5. The level of significance was $\alpha < 0.05$.

**Detailed Results**

*Comparisons between mPFC and vHPC: the role of optogenetic stimulation.* At first, we evaluated if the animal groups learned the initial place avoidance task equally. A two-way RM ANOVA revealed an effect of the days (Acq1, Acq2, Acq3) in the n° of entrances (F [1.959, 54.84] = 29.92; p<0.0001), in the latency to target sector (time to first entrance) (F [1.675, 46.91] = 24.71; p<0.0001), and in the time spent in the target sector (F [1.745, 48.87] = 8.542; p=0.0011) suggesting that mice of all groups learned the initial place avoidance task equally.

Subsequently, we evaluated whether the mice could reverse the previous spatial acquisition based on the RF rule. A two-way RM ANOVA revealed an effect of the days (Rev1, Rev2, Rev3) in the time spent in the target sector (F [1.966, 55.04] = 16.16; p<0.0001), suggesting that optogenetic stimulations in both mPFC and vHPC at specific frequencies helped to reverse the previously acquired target sector. Furthermore, an effect of the groups was found in the total distance moved (F [3, 28] = 11.38; p<0.0001) and in the latency to the target sector (F [3, 28] = 15.54; p<0.0001). Finally, multiple comparisons confirmed the rescuing ability of the optogenetic stimulations in mPFC (50Hz) to their WTs and the stimulations in vHPC (10Hz).

Following, we evaluated whether the mice could flexibly shift the RF-to-AF rule and navigate flexibly in the arena. A two-way RM ANOVA revealed an effect of the groups in the total distance moved (F [3, 28] = 11.56; p<0.0001), in the n° of entrances (F [3, 28] = 13.67; p<0.0001), in the latency to target sector (F [3, 28] = 7.089; p=0.0011), and in the time spent in target sector (F [3, 28] = 4.521; p=0.0105) suggesting that optogenetic stimulations in the mPFC at 50Hz significantly helped in flexibly shifting the RF-to AF rule to their WTs, and significantly better helped the flexibility in respect with the vHPC PV-Cre group. Moreover, an effect of the days was found in the latency to the target sector (F [1.912, 53.53] = 3.846; p=0.0292) and in the time spent in the target sector (F [1.885, 52.77] = 5.515; p=0.0076). Finally, multiple comparisons suggest an overall ability to switch from reversal spatial memory to navigational flexibility when mPFC and vHPC are stimulated at specific frequencies.

However, the previous analysis only established that optogenetic stimulations in both mPFC and vHPC faced the MK801 challenge in reversal spatial memory and navigational set-shifting during the days. These results did not fully clarify if PV+ optogenetic stimulations in mPFC and vHPC could rescue those executive functions. To verify this, we compared the last day of acquisition (Acq3) with the reversal (Rev1, Rev2, Rev3) and the last day of reversal (Rev3) with the set-shifting (Set1, Set2, Set3). Analysis was performed using two-way RM ANOVAs on the previously measured behavioral parameters. Results are shown in Table 3 (Tab. 3). Two-way RM ANOVA comparing Acq3 with reversal revealed a significant effect of the sessions in the n° of entrances (F [2.036, 63.10] = 24.56; p<0.0001) in the latency to the target sector (F [1.335, 41.39] = 45.40; p<0.0001), and the time spent in the target sector (F [2.719, 76.12] = 41.17; p<0.0001). Furtherly, a significant effect of the groups was found in the total distance moved (F [3, 28] = 11.77; p<0.0001), in the latency to the target sector (F [3, 28] = 18; p<0.0001), and in the time spent in the target sector (F [3, 28] = 16.49; p<0.0001). Multiple comparisons showed that $\gamma$-like 50Hz optogenetic stimulations in the mPFC had an immediate and generally lasting effect in reversing the spatial memory of the to-be-avoided sector compared to the WT and the PV-Cre-vHPC groups. Of note, the vHPC-PV-Cre group showed better ability to reverse the time spent in the target sector than the mPFC-Pv-Cre group, suggesting that $\theta$-like 10Hz optogenetic stimulations rescued the flexibility to reverse the previous spatial memory better than $\gamma$-like 50Hz in mPFC.

Two-way RM ANOVA comparing Rev3 with set-shifting revealed a significant effect of the groups in the total distance moved (F [3, 28] = 10.18; p=0.0001), n° of entrances (F [3, 28] =

14.21; p<0.0001), in the latency to the target sector (F [3, 28] = 7.482; p<0.0001), and the time spent in the target sector (F [3, 28] = 10.46; p<0.0001). Also, a significant effect of the sessions was found in the total distance moved (F [2.709, 75.85] = 9.474; p<0.0001), in the n° of entrances (F [2.419, 67.73] = 5.921; p=0.0025), in the latency to the target sector (F [2.588, 72.48] = 5.56; p=0.0020), and in the time spent in the target sector (F [2.797, 78.31] = 7.362; p=0.0003). Multiple comparisons suggest that $\gamma$-like 50Hz optogenetic stimulations in the mPFC had an immediate and generally lasting effect in flexibly shifting the RF-to-AF navigation rule compared to the WT and PV-Cre-vHPC groups. However, the vHPC-PV-Cre group showed a better ability to shift the RF-to-AF navigation rule, spending less time in the target sector than the mPFC-Pv-Cre group. Conversely, the mPFC-PV-Cre group showed a better ability to "re-learn" the new navigation rule during the set-shifting days than the vHPC-PV-Cre group.

*Comparisons between optogenetic stimulations and pharmacological control groups (NaCl vs. MK801): a resemblance with the NaCl groups.* Results are summarized in Table 4 (Tab. 4). A two-way RM ANOVA on reversal revealed an effect of the groups in the total distance moved (F [5, 38] = 45.13; p<0.0001), n° of entrances (F [5, 38] = 33.09; p<0.0001), in the latency to the target sector (F [5, 38] = 24.23; p<0.0001), and the time spent in the target sector (F [5, 38] = 10.28; p<0.0001), suggesting that WT-mPFC group resembled MK801 group, and that mPFC-PV-Cre and NaCl behaved similarly. Furtherly, it has been found an effect of the days in the n° of entrances (F [1.726, 65.59] = 84.53; p<0.0001) and in the time spent in the target sector (F [1.878, 71.38] = 20.78; p<0.0001) suggesting that both PV-Cre groups (mPFC and vHPC) and NaCl group showed an ability to reverse the previous spatial memory throughout the days. A two-way RM ANOVA on set-shifting revealed an effect of the groups in the total distance moved (F [5, 38] = 18.95; p<0.0001), the n° of entrances (F [5, 38] = 15.69; p<0.0001), in the latency to the target sector (F [5, 38] = 10.76; p<0.0001), and the time spent in the target sector (F [5, 38] = 7.474; p<0.0001). Multiple comparisons suggested that $\theta$-like 10Hz optogenetic stimulations in the PV-Cre vHPC group did not produce any overall resemblance to the NaCl group. However, as previously mentioned, PV-Cre vHPC showed similarities with the NaCl group for the time spent in the target sector – a measure of flexible shifting of the RF-to-AF navigation rule. Furtherly, it has been found an effect of the days in the n° of entrances (F [1.701, 64.62] = 9.301; p=0.0006), in the latency to the target sector (F [1.941, 73.76] = 11.66; p<0.0001), and in the time spent in the target sector (F [1.815, 68.97] = 6.302; p<0.0001). Multiple comparisons in the time spent in the target sector suggested a general ability to learn the new navigation rule in WT-mPFC and MK801 groups. At the same time, the PV-Cre groups (mPFC and vHPC) and NaCl group were not able to increase the shifting ability throughout the days, furtherly suggesting that both $\gamma/\theta$-like optogenetic stimulations induced similar cognitive abilities to the naïve NaCl group.

*Comparisons with NaCl control groups: ruling out potential effects of MK801 systemic injection.* As previously mentioned, all the experimental and the control groups received NaCl instead of MK801 systemic injections and optogenetic stimulations during the reversal and the set-shifting days. A two-way RM ANOVA on reversal revealed an effect of the groups in the total distance moved (F [3, 20] = 11.29; p=0.0002), the n° of entrances (F [3, 20] = 8.334; p=0.0009), in the latency to the target sector (F [3, 20] = 4.929; p=0.0101), and in the time spent in the target sector (F [3, 20] = p=0.0010) suggesting that NaCl systemic injections-optogenetic stimulations worsen the inflexibility, especially in the vHPC groups. An effect of the days was found in the latency to enter (F [1.616, 32.31] = 8.535; p=0.0020) and the time spent in the target sector (F [1.788, 35.76] = 14.71; p<0.0001), thus confirming that combinations of NaCl i.p. and optogenetic stimulations of PV+ interneurons were ineffective in rescuing reversal spatial memory. Subsequently, a two-way RM ANOVA on set-shifting revealed an effect of the groups in the total distance moved (F [3, 20] = 7.177; p=0.0019), the n° of entrances (F [3, 20] = 8.737; p=0.0007), in the latency to the target sector (F [3, 20] = 24.84; p<0.0001), and in the time spent in the target sector (F [3, 20] = 12.11; p<0.0001) confirming that MK801-mPFC group performed better than its NaCl group in all the behavioral parameters. These results suggest that the combination of NaCl systemic administration and optogenetic stimulation of PV+ interneurons in mpFC and vHPC exacerbate navigational inflexibility.

*Effectiveness of the viral transfections and confirmation of the local optogenetic stimulations.* *Effectiveness of the viral transfections and confirmation of the local optogenetic stimulations.* The primary goals of these analyses were to confirm the effectiveness of the local viral transfections of ChR2 on PV+ interneurons in both mPFC and vHPC and to confirm that optogenetic stimulations induced behavioral flexibility during the reversal and the set-shifting sessions (Fig. 6). Firstly, we counted the PV+ cells in the region of interest (ROI, $10mm^2$) as % of the fluorescent area (% area) at rostro-caudal layers (+2.1 to +1.3 AP with respect to bregma, mPFC; -2.7 to -3.5 with respect to bregma, vHPC). A RM one-way ANOVA found a significant effect of optogenetic stimulations in mPFC (F [1.178, 17.67] = 29.69; p<0.0001) (Fig 6a), and vHPC (F [1.357, 20.35] = 40.94; p<0.0001) (Fig. 6b). Comparisons showed a significant difference between the caudal and rostral slices and the optic fiber position in mPFC (F [15, 45] = 4.940; p<0.0001) and vHPC (F [15, 45] = 4,534; p<0.0001). Further, we analyzed the percentage of the area (% area) (Fig. 6c) and the integrated density (integrated density) of fluorescence (area x mean gray value) for ChR2/PV+ interneurons and c-Fos protein expressions (Fig. 6d). For the % area, a one-way ANOVA showed an effect of the optogenetic stimulations (F [7, 128] = 10,54; p<0.0001). Multiple comparisons interestingly showed that mPFC-WT group had a significative increase of c-Fos expression despite the ineffectiveness of PV+ optogenetic stimulations (p=0.0084). Also, a significative difference between the expression of PV+/ChR2 and c-Fos protein in the vHPC-PV-Cre group (p<0.0001) was found, suggesting that despite the optogenetic stimulations, PV+ were not activated by those stimulations (Fig. 6c). A one-way ANOVA for the integrated density substantially replicated the previous results for the % of the area (F [7, 128] = 15,61; p<0.0001). Multiple comparisons confirmed a significative difference between the expression of PV+/ChR2 and c-Fos protein in the vHPC-PV-Cre group (p<0.0001), confirming that despite the optogenetic stimulations, PV+ were not activated by those stimulations (Fig. 6d). Of note, the expression of c-Fos protein in the WT groups of both mPFC and vHPC was similar to that of PV-Cre groups, ruling out potential adverse effects of optogenetic stimulations. Moreover, we analyzed the level of colocalization of ChR2/PV+ interneurons and c-Fos protein around the tip of the optic fiber (ROI, $1mm^2$). A one-way ANOVA analysis of colocalization between PV+/ChR2 and c-Fos protein showed an extreme significance in both PV-Cre groups (F [3, 64] = 105.6; p<0.0001), suggesting that optogenetic stimulations indeed activated PV+ interneurons (Fig. 6e). In Figure 6e and 6f, representative areas and cells of mPFC and vHPC, respectively, are shown.
